# Sex-biased transcriptomic response of the reproductive axis to stress

**DOI:** 10.1101/152801

**Authors:** Rebecca Calisi, Suzanne H. Austin, Andrew S. Lang, Matthew D. MacManes

## Abstract

Stress is a well-known cause of reproductive dysfunction in many species, including birds, rodents, and humans. However, little is known of the genomic basis for this dysfunction and how it may differ between the sexes. Using the classic reproductive model of the rock dove (*Columba livia*), we conducted the most in-depth investigation to date of how stress affects all gene transcription of a biological system essential for facilitating reproduction - the hypothalamic-pituitary-gonadal (HPG) axis. The HPG transcriptome responded to stress in both sexes, but females exhibited more differential expression than males, and these stress responsive genes were mostly unique to females. This result may be due to 1) fluctuations in the female endocrine environment to facilitate ovulation and follicle maturation, and 2) their evolutionary history. We offer a vital genomic foundation on which sex-specific reproductive dysfunction can be studied, as well as novel gene targets for genetic intervention and therapy investigations.

## INTRODUCTION

Stress can disrupt reproduction in multiple, complex ways [1–3]. The perception of a stressor activates the hypothalamic-pituitary-adrenal (HPA) axis, which results in a synthesis of stress hormones (glucocorticoids) [4]. Glucocorticoid hormones (cortisol in humans, corticosterone in birds and rodents) are synthesized by the adrenal cortex and exert both rapid and gradual actions on vertebrate physiology [5]. This activation of the HPA system can cause suppression of the reproductive system, i.e., the hypothalamic-pituitary-gonadal (HPG) axis, at multiple levels (Fig. 1), including inhibiting gonadotropin-releasing hormone (GnRH) secretion from the hypothalamus, suppressing luteinizing hormone (LH) and follicle stimulating hormone (FSH) release from the pituitary, sex steroid hormone release from the gonads, and ultimately reducing or eliminating sexual behavior and reproduction [6–9]. However, questions remain as to 1) how stress affects all gene activity of the HPG axis, and 2) if these effects are sex-specific. Evidence suggests regulatory mechanisms of the HPG system under stress can be sex-specific (eg. human: [10]; rodent: [11]; birds: [12,13]), but the full extent of sex-biased changes is still largely unknown. In general, males have historically dominated animal studies [14–16], obscuring discovery of potential sex differences that could inform and guide further research and clinical studies [17].

**Figure 1.**
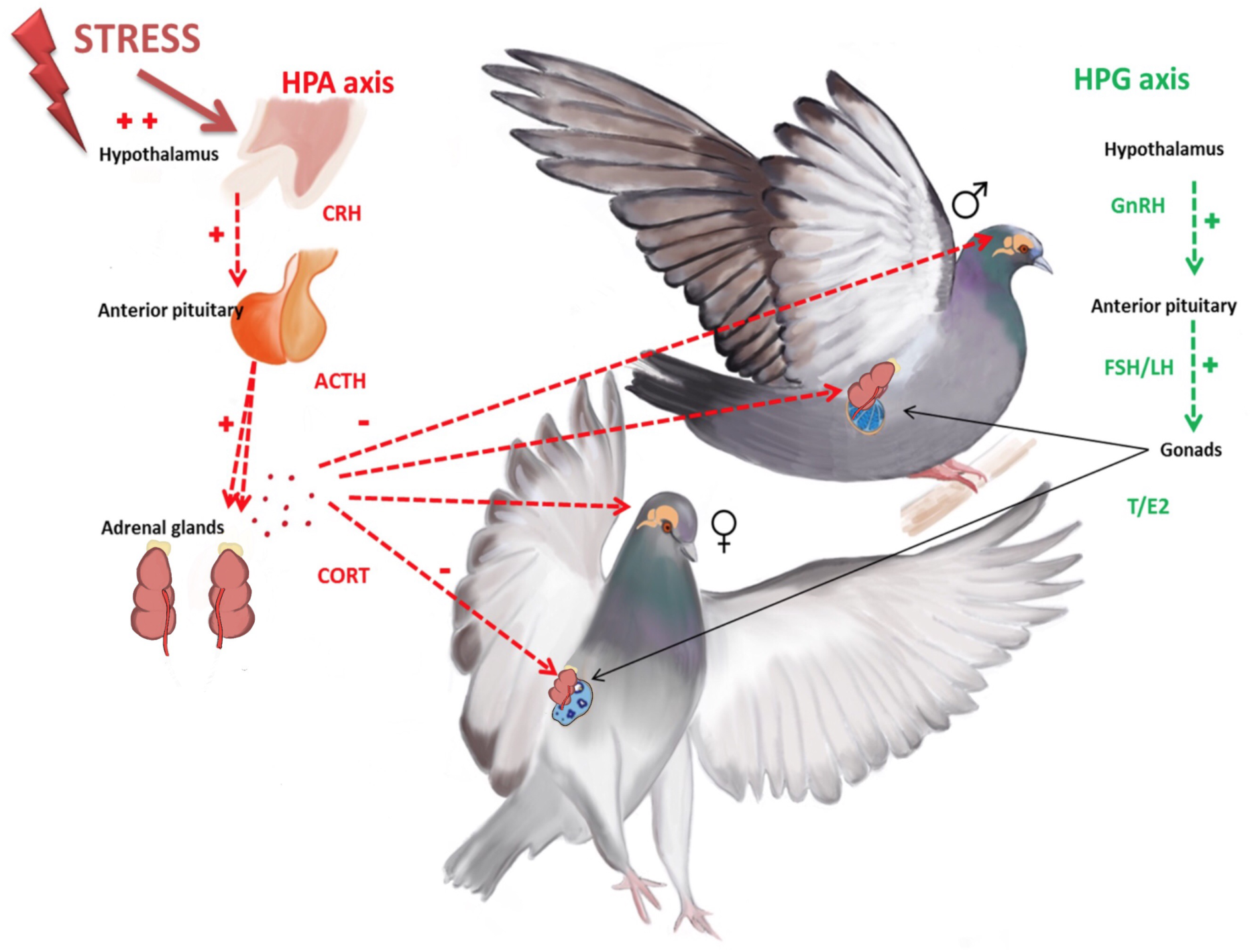
Depiction of the hypothalamic-pituitary-adrenal (HPA), or “stress”, axis and its intersection with the hypothalamic-pituitary-gonadal, or “reproductive”, axis (HPG). Illustration by Natalia Duque.

Here, we tested the effects of stress on the genomic activity of the male versus female HPG axis using the model of the rock dove (*Columba livia*). Doves have been historically used to study reproductive behavior [18–20] and now are proving to be a valuable model for genomics research [21–24]. We exposed sexually mature males and females to 30 min of restraint stress, which successfully activates the stress response as measured through significantly increased circulating plasma glucocorticoids. We compared the genomic expression of the HPG axis of stressed females and males to each other and in comparison with unstressed controls. We report patterns of tissue-specific and sexually dimorphic gene expression, with females showing a greater stress response at the level of the transcriptome in all three tissues - the hypothalamus, pituitary, and gonads - as compared to males. These data offer a valuable resource to advance stress and reproductive research with the potential for devising future therapeutic strategies to ameliorate stress-induced HPG axis dysfunction.

## RESULTS

### Corticosterone Assay

Plasma corticosterone was assayed from 48 birds (stress treatment group: 12 male, 12 female; control group: 12 male, 12 female) using a rodent Corticosterone RIA kit and a 1:20 dilution (MP Biomedicals, Orangeburg, NY). Plasma corticosterone concentrations were significantly higher in both stressed birds as compared to controls (treatment: F_1,46_=73.5, P<0.001). Neither sex nor an interaction effect of treatment*sex were statistically significant (sex: F_1, 46_=0.5, P=0.505; treatment* sex=F_1,46_=1.0, P=0.333).

### Sequence Read Data & Code Availability

In total, 24 hypothalami (12 male, 12 female), 24 pituitary glands (12 male, 12 female), 12 testes, and 12 ovaries from 24 birds were sequenced. Each sample was sequenced with between 2.3 million and 24.5 million read pairs. Read data corresponding to the control birds are available using the European Nucleotide Archive project ID PRJEB16136; read data corresponding to the stressed birds are available at PRJEB21082. Code used for the analyses of these data are available at https://git.io/vPA09.

### Transcriptome assembly characterization

The Rock Dove version 1.1.0 transcriptome (available https://goo.gl/S8goSM, and at Dryad *post acceptance*) contains 92,805 transcripts, of which 4,794 were added as part of this study to the previous version 1.0.4 transcriptome. This newly compiled transcriptome data improves genic contiguity, increasing the number of complete BUSCOs 0.2% to achieve 86.1% relative to the version 1.0.4 assembly.

### Sequence Read Mapping and Estimation of Gene Expression

Raw sequencing reads corresponding to individual samples of hypothalami, pituitary glands, and gonads were mapped to the Rock Dove reference HPG axis transcriptome version 1.1.0 using Salmon, resulting in 80% to 90% read mapping. These data were imported into R and summarized into gene-level counts using tximport, after which, edgeR was used to generate normalized estimates of gene expression. 17,263 genes were expressed in the HPG.

### Evaluation of Candidate Gene Expression

Using the assembled transcriptome, HPG-specific and sexually dimorphic gene expression patterns were characterized in stressed versus control animals. *A priori*, target genes were identified for investigation that are well known to play a role in reproduction and the stress response (Table 1). Using a generalized linear model and least-squares means for post-hoc tests of significance (P<0.05), we found sex and tissue-specific differences in HPG transcriptomic activity in response to the stress treatment.

**Table 1.**
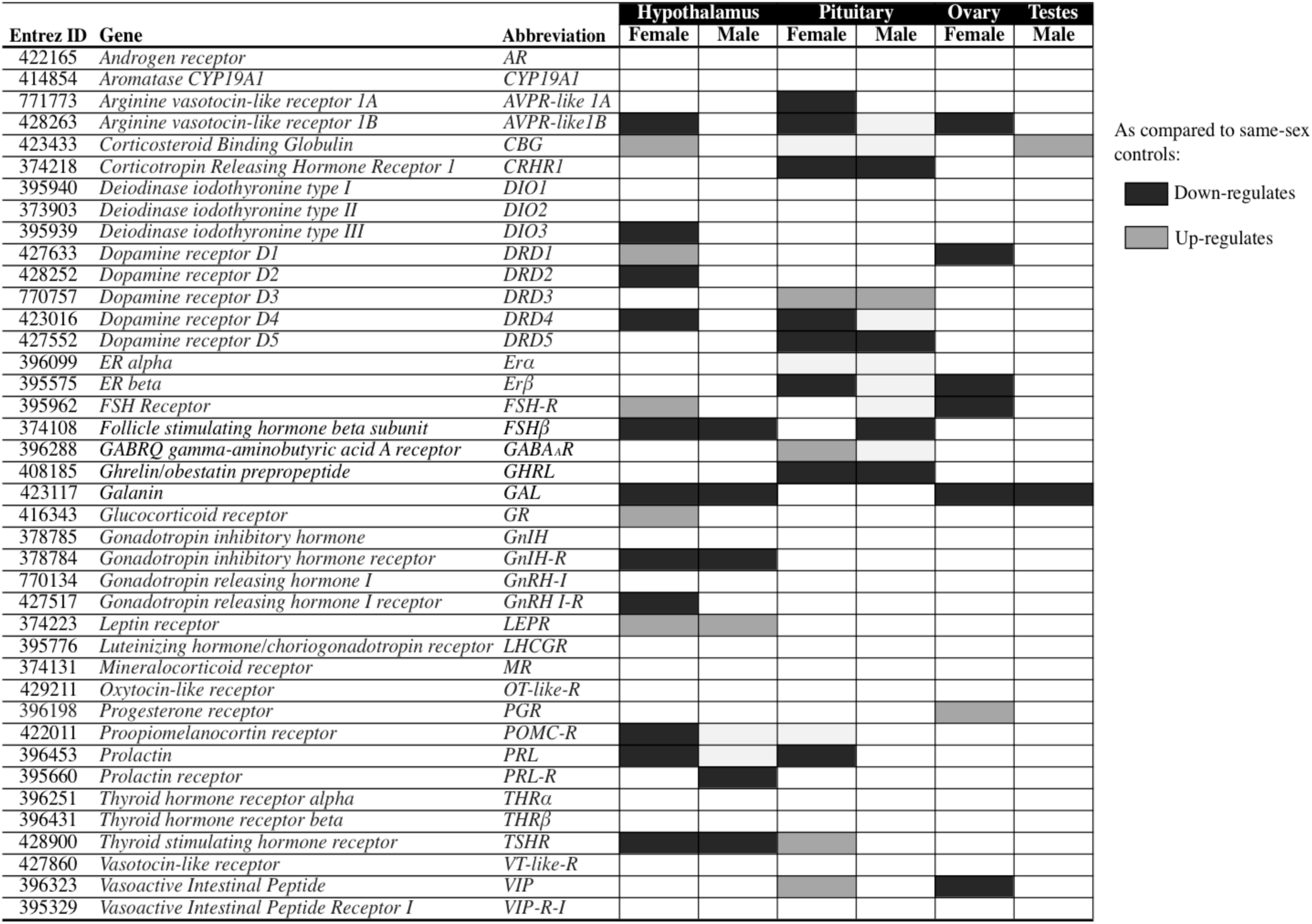
Results of candidate gene expression analysis. Target genes identified *a priori* due to their known role in reproduction and/or the stress response. Expression of these genes was present in all HPG axis tissues. Genes that significantly (P<0.05) upregulated in expression in response to stress are indicated with a lighter shade; genes that significantly dowregulated in response are indicated with a darker shade.

### Global Evaluation of Gene Expression

Global patterns of gene expression were analyzed using edgeR to observe how the entire transcriptome of the HPG axis responds to stress. After controlling for over 17,000 multiple comparisons, the count data was normalized using the TMM method [25], which, in brief, uses a set of scaling factors for library sizes to minimize inter-sample log-fold changes for most genes. This analysis revealed a significant transcriptomic response of the HPG axis to stress, particularly in the female, especially in the pituitary and the ovary (Table 2, Fig. 2, 3). A complete list of differentially expressed genes in female and male HPG tissue in response to stress can be found at https://goo.gl/mypuFv. A brief description of each gene’s reported functionality in vertebrates is given in forthcoming text to offer insight into potential systems affected by stress. However, it is important to note that the gene functionality given may not have yet been confirmed in an avian model.

**Figure 2.**
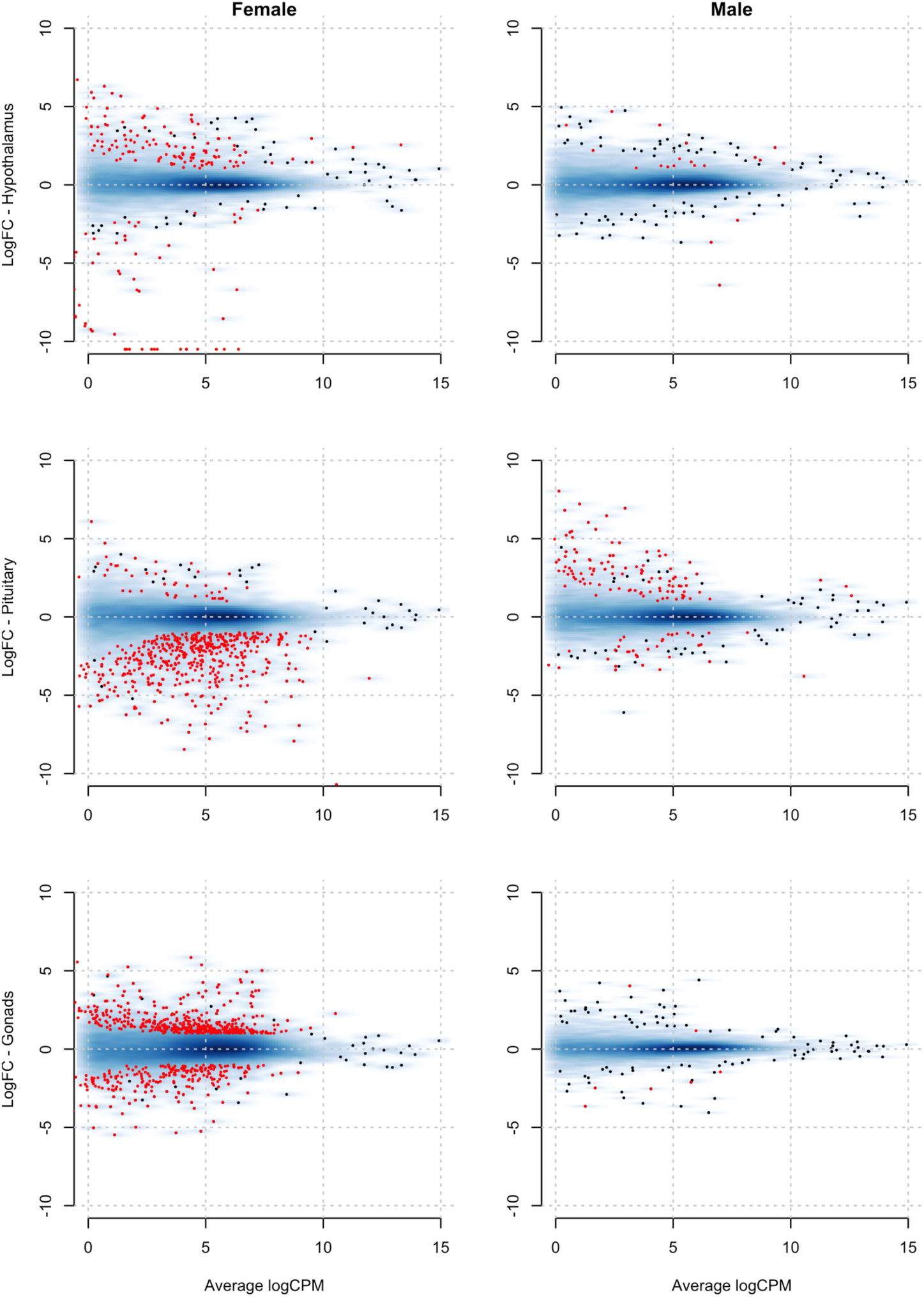
MA plots depicting the amount of genes differentially expressed throughout the HPG axis of females (left column) as compared to males (right column). Red dots indicate statistically differentially expressed genes. The X axis represents the average value for gene expression (units = average counts per million). The Y axis indicates the log fold change (LogFC) of genes expressed in the hypothalamus (top), pituitary (middle) and gonads (bottom). Red dots above zero indicate higher expression in the control animals; red dots below zero indicate higher expression in stressed animals.

**Fig. 3.**
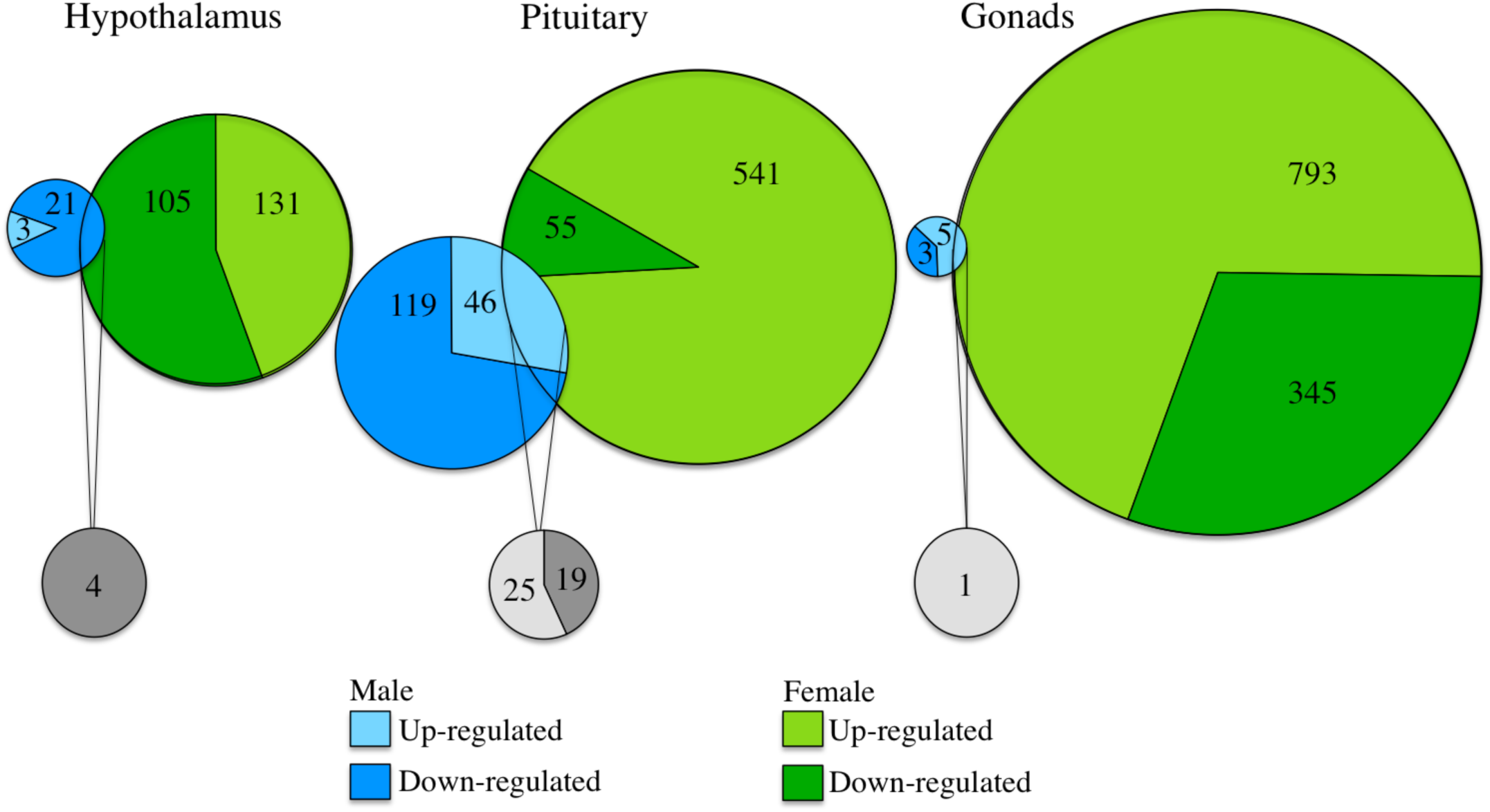
A weighted Venn diagram depicting the overlap of the number of differentially expressed genes between the sexes in the hypothalamus, pituitary, and gonads in response to stress as compared to controls. Genes that upregulated in expression in response to stress are shown in a lighter color; genes that dowregulated in response exhibit a darker color. Numbers within shaded areas indicate the number of stress-responsive genes. Pie charts of shared, stress-responsive genes have been magnified and are not to scale.

**Table 2.** The number of differentially expressed (DE) genes in each tissue in response to stress as compared to controls.

| Tissue | Sex | In response to stress: |  |  |
| --- | --- | --- | --- | --- |
|  |  | Higher DE | Lower DE | Total DE |
| Hypothalamus | Female | 105 | 131 | 136 |
|  | Male | 3 | 21 | 24 |
| Pituitary | Female | 541 | 55 | 596 |
|  | Male | 46 | 119 | 165 |
| Ovary | Female | 345 | 793 | 1,138 |
| Testes | Male | 5 | 3 | 8 |

### Hypothalamus

A global transcriptome analysis yielded 236 differentially expressed genes in the hypothalami of stressed females (106 upregulated and 131 downregulated) as compared to female controls, while 24 genes were significantly differentially expressed in the hypothalami of stressed males (3 upregulated and 21 downregulated; Table 2, Fig. 3) as compared to male controls. Genes more highly expressed in the hypothalamic tissue of stressed females include *UNC45B*, which is a progesterone regulator and co-chaperone for heat shock protein of 90 kDa (*HSP90*) [26] (logFC -8.6, FDR 9.0e-12). Additionally, *interferon induced transmembrane protein 5* (*IFITM5*), which was more highly expressed, can interact with serotonin receptor [27] as well as several solute carriers [28] responsible for transporting newly synthesized prostaglandins PGD2, PGE1, PGE2, leukotriene C4, and thromboxane B2 (logFC -11.1, FDR 3.0e-10). Several long non-coding RNAs, *LOC107050862* and *LOC101747554* were some of the most differentially expressed genes between stress and control groups (logFC -12.6, FDR 1.0e-13 and logFC -11.3, FDR 4.5e-11 respectively), yet their function is currently unknown. In addition to these, a strong signal of differential expression of the Myosins was uncovered (*e.g., MYH1E, MYH1F, MYOZ2*). Gene ontology terms enriched for this treatment group to describe groups of genes of similar function responsive to stress include terms related to muscle development and function (gene ontology term myofibril assembly, GO:0030239). This pattern is driven in part by Myosin proteins, which can play a role in secretory function [29,30].

Genes more lowly expressed in the hypothalamic tissue of stressed females as compared to controls include *LUC7L2* (logFC 1.9, FDR 1.1e-07), a gene previously found to be differentially expressed in response to stress [31], and *LOC107051530* (logFC 4.9, FDR 6.4e-5), a non-coding RNA whose function is unknown. *Prolactin* is more lowly expressed in response to stress (logFC 2.5, FDR .0002), as is *GABA type A receptor-associated protein* (logFC 1.2, FDR .007), the latter which has been shown to be regulated by *Leptin* in guinea pigs [32]. No significantly enriched gene ontology terms were revealed for genes more lowly expressed in the female hypothalamus in response to stress. The full differential dataset for the female hypothalamus is available at https://git.io/vD4LU.

As compared to the females, fewer genes were differentially expressed in the male hypothalamus in response to stress. One gene more highly expressed in stressed males as compared to controls is *PH domain leucine-rich repeat-containing protein phosphatase 2* (*PHLPP2*), which can play a role in the apoptotic process (logFC-3.6, FDR 2.4e-09). A gene more lowly expressed in response to stress, *Pro melanin concentrating hormone* (*PMCH*) (logFC 3.8, FDR 2.532022e-05), can inhibit stress-induced ACTH release, stimulate anxiety and sexual behavior, and antagonize the inhibitory effect of alpha melanotropin on exploration behavior [33–35]. *C-RFamide protein*, also known as *prolactin releasing hormone 2* [36] was also more lowly expressed in response to stress and can play a role in the release of prolactin as well as several gonadotropes [37] (logFC 4.7, FDR 2.52e-05). No significantly enriched gene ontology terms were revealed for genes differentially expressed in the male hypothalamus in response to stress The full differential dataset for the stress response of the male hypothalamus is available at https://git.io/vD40k.

### Pituitary Gland

A global transcriptome analysis of the HPG stress response yielded 596 differentially expressed genes in the pituitary of stressed females (541 upregulated and 55 downregulated) as compared to female controls, while 165 genes were differentially expressed in the pituitary of stressed males (46 upregulated and 119 downregulated; Table 2, Fig. 3) as compared to male controls. For example, in females, *Proteolipid Protein 1* (*PLP1* - [38]), a gene implicated in the stress response, as well as *Myelin basic protein* (*MBP)* [39], which can interact with PLP1 to regulate apoptosis, are more highly expressed in response to stress (logFC -10.7, FDR 4.4e-12 and logFC -3.9, FDR 4.4e-12). Genes like *Cytokine inducible SH2 containing protein* (*CISH*) and *glutamate metabotropic receptor 3* (*GRM3*) are also more highly expressed in females in response to stress (logFC -1.8, FDR 1.9e-10; logFC -3.8, FDR 4.2e-10, respectively). *CISH* is involved in the immune response via negatively regulation of cytokine signalling [40], and *GRM3* has been reported as a candidate gene for schizophrenia [41]. Gene ontology terms enriched for genes more highly expressed in the female pituitary in response to stress are related to oligodendrocyte differentiation, positive regulation of gliogenesis, G-protein coupled receptor signaling pathway, and behavior.

Genes more lowly expressed in the pituitary of stressed females as compared to controls include *Angiopoietin like 7* (*ANGPTL7*) (logFC 2.6, FDR 5.9e-5) whose function is currently unknown but may be related to the regulation of growth and metabolism (Hato et al. 2008). *Chemokine* (*CC motif*) *ligand 4* (*CCL4*), a gene previously implicated in the heat stress response in chickens (Tu et al. 2016), is also more lowly expressed in response to stress. No significantly enriched gene ontology terms were revealed for genes differentially expressed in the female pituitary in response to stress. The full differential dataset for the stress response of the female pituitary is available at https://git.io/vD49n.

Much like the pattern observed in the hypothalami of males as compared to females, the male pituitary also has less differential gene expression than the female pituitary. For example, genes that were more highly expressed in the male pituitary in response to stress include *Nuclear receptor subfamily 4 group A member 3* (*NR4A3*) (logFC -2.9, FDR 5.3e-10), which has been implicated in mechanisms related to feeding behavior and energy balance [42,43], and 5*-hydroxytryptamine receptor 3A* (*HTR3A*) (logFC -1.9, 4.42e-05), a receptor whose ligand (serotonin) is widely accepted to be involved in mood disorder and anxiety [43–46]. A signal of cell-cycle arrest may be evident, as the gene *CCAAT/enhancer binding protein* (*C/EBP*), *delta*, know to be involved in preventing the cell cycle from continuing through the G1 phase [47] is more highly expressed in response to stress (logFC -2.6, FDR 3.1e-09). *Activating transcription factor 3* (ATF3), which has been previously reported to respond to stress [48,49], is highly expressed in the male pituitary in response to stress. Gene ontology terms enriched for in genes more highly expressed in the stressed male pituitary in response to stress are related to the term for “muscle development”. Again, this signal is descendent from actin and myosin genes involved in secretion [29] as opposed to actual skeletal or smooth muscles.

Genes more lowly expressed in the male pituitary in response to stress include the *class II major histocompatibility complex (MHC) antigen and DM beta chain type 1* (DMB1) (logFC 2.11 FDR 0.0009), which has important implications for stress given their well known roles in immune function [50,51]. Other genes include the Heat shock 27kDa protein 1 (HSPB1) (logFC 2.8., FDR 0.008) and Synaptotagmin 2 (logFC 1.21, FDR 0.009), the latter which is know to be involved in the stress response. [52,53]. The full table of differential expression results for the male pituitary in response to stress is available at https://git.io/vD4xL.

### Ovary

The ovary was the site of the most differential expression in response to stress as compared to the hypothalamus, pituitary, and testes. In sum, 1138 genes were differentially expressed (345 higher in stress, 793 lower; Table 2, Fig. 3) in the ovary of stressed females as compared to controls. Genes that were more highly expressed include Heat Shock Protein 25 (HSP25) (logFC -2.9, FDR 5e-07) and *Heat shock 70kDa protein 8* (*HSPA8*), logFC -1.5, FDR 2.5e-08), the latter has been reported to prevent protein aggregation under stress conditions [54]. Additionally, *Thyroid hormone responsive protein* (*THRSP*) was more highly expressed in the ovary in response to stress (logFC -5.1, FDR 1.1e-05). *THRSP* has been reported to play a role in the regulation of lipogenesis, especially in lactating mammary gland [55]. Gene ontology terms enriched for genes more highly expressed in the ovary in response to stress are related to inflammatory response, defense response, and response to stress.

### Testes

Analysis of differential expression in the testes in response to stress resulted in only 8 differentially expressed genes (Table 2, Fig. 3). These genes include *Heat shock 27kDa protein 1* (*HSPB1*) (logFC -2.5, FDR 0.005) and *Thyroid hormone receptor interactor 11* (*TRIP11*) (logFC -1.55, FDR 0.01), both of which increase in expression in response to stress as compared to controls. *TRIP11* has been previously implicated in the stress response [56], yet its function in the testes is currently unknown. The full table of differential expression results for the male testes in response to stress is available at https://git.io/vDBTw.

### Sex-biased Response to Stress

In the previous analyses, changes in gene expression in response to restraint stress were identified in each sex as compared to same-sex controls. Here, changes in gene expression in response to restraint stress are compared between the sexes, specifically in hypothalamic and pituitary tissue. Male and female gonads are structurally and functionally different and thus were not directly compared. A distributional analysis was used to identify genes whose expression patterns varied by sex. Specifically, the sex difference in gene expression in response to stress (defined as (Median Expression _Male Tissue Stress_ - Median Expression _Male Tissue Control_) -(Median Expression _Female Tissue Stress_ - Median Expression _Female Tissue Control_) was calculated, with genes whose response was different (defined as being more responsive than 98% of all other genes, n=173 in each tail of the distribution, Fig. 4), deemed to be expressed in a sex-biased manner.

**Figure 4.**
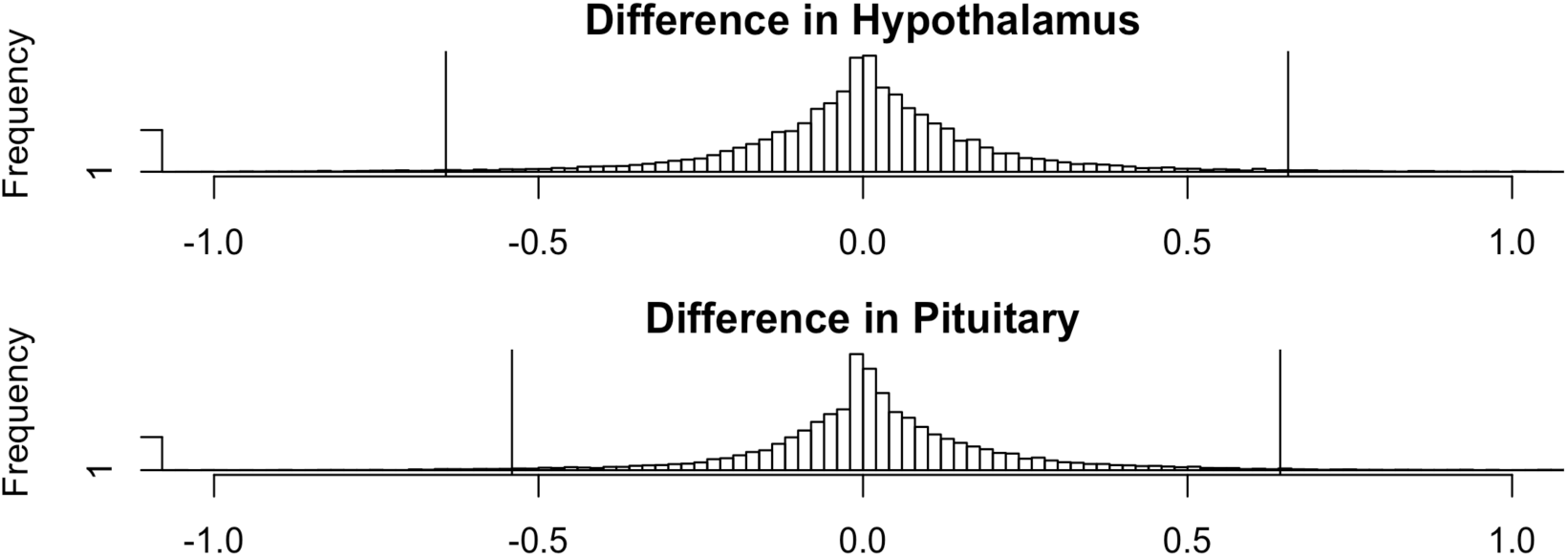
The transcriptomic stress response distribution histogram in males versus females in the hypothalamus (top) and pituitary (bottom). Vertical lines in the tails of the distributions indicate genes in the upper and lower 1% of the distribution, beyond which genes were deemed to be expressed in a sex-biased manner. The X axis represents the difference in expression response, and the Y axis represents the frequency. To the right of 0.0 point are genes more highly expressed in males, while genes to the left of 0.0 point are more highly expressed in females.

### Hypothalamus

Genes that were more responsive to stress in the female hypothalamus relative to the male hypothalamus (defined as ≥ 99% differences; examples in Figure 5a) included *Cholecystokinin* (*CCK;* ΔMedian= 0.75), which has been previously implicated in opiate antagonism [57], appetite [58], and the stress response [59,60]. Another gene of interest discovered to be more responsive in females was *Calcitonin Gene-Related Peptide I* (*CALCA*, also known as CGRP, ΔMedian= 1.02), known to be an important regulator of stress-induced reproductive suppression [61]. Actin beta-like 2 (ACTBL2), a gene whose function may be linked to secretory function [29] as well as to leptin and insulin mediated signalling [62], were more responsive to stress in the female hypothalamus as compared to the male hypothalamus (ΔMedian = 1.39), as was *Growth Hormone* (GH) (ΔMedian = .77), which can stimulate growth and cell reproduction as well as respond to stress by increasing glucose and fatty acids. The full table of genes, more responsive to stress in the female hypothalamus as compared to the male hypothalamus is available at https://git.io/vDzKT.

**Figure 5.**
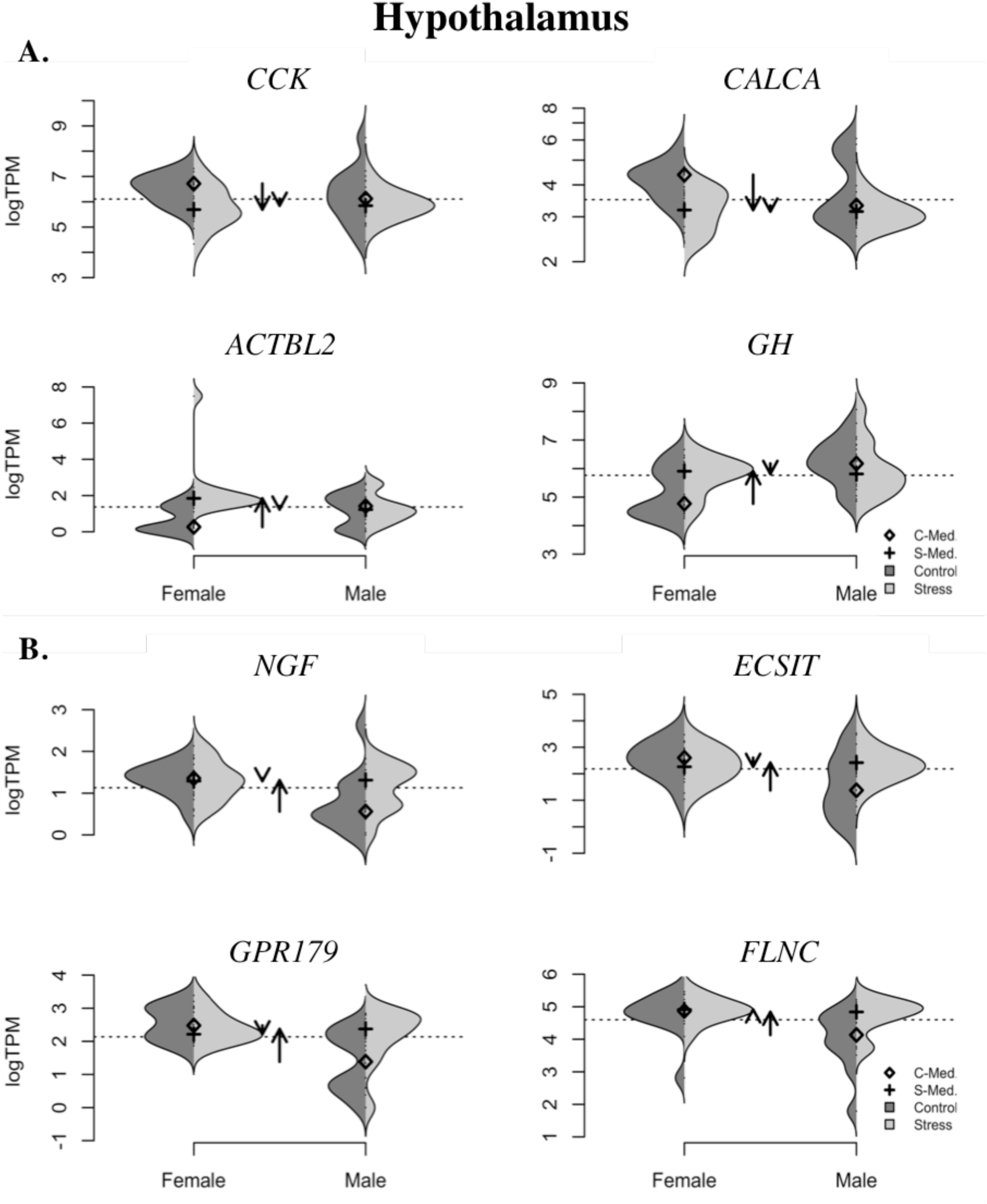
Violin plots depicting genes that were more responsive to stress in the a) female hypothalamus relative to the male hypothalamus, and b) male hypothalamus relative to the female hypothalamus. Plot titles are of the format Gene Name:Entrez ID:Tissue. The Y-axis indicates levels of gene expression as presented by taking the log of transcripts per million (logTPM). The shape of each half of the plot represents the kernel density estimation of the data, with darker grey indicative of data from control animals and lighter grey from stressed animals. The diamond symbol represents the median value of the control group while the plus sign represents the median value of the stressed group. Arrows represent the direction and amount of change in gene expression in the stressed group as compared to controls (up for upregulated, down for downregulated).

Genes that were more responsive to stress in the male hypothalamus relative to the female hypothalamus (examples in Figure 5b) included Toll-like Receptor 15 (TLR15) and MHC DMbeta2 (DMB2), which play a role in immune function (ΔMedian= 0.74 and 0.89, respectively)[63,64] which can affect the stress response [65]. Hypothalamic genes more highly responsive to stress in males versus females appear to be related to Mitogen-Activated Protein Kinase (MAPK) activity, including nerve growth factor (NGF), Filamin C (FLNC), and ECSIT Signalling Integrator (ECSIT). The MAPK cascade is thought to coordinate the response to a wide variety of stressors, given it receives stimuli from a diverse group of signalling pathways including growth factors, G protein-coupled receptors, pathogen-associated molecular patterns (PAMPs) and danger-associated molecular patterns (DAMPs), reviewed in [66]. The full table of differential expression results of genes more responsive to stress in the male hypothalamus as compared to the female hypothalamus is available at https://git.io/vDz9C.

### Pituitary

Genes more responsive to stress in the female pituitary relative to the male pituitary include (examples in Fig. 6a) melanin concentrating hormone receptor 1 (MCHR1), ΔMedian= 1.32), a gene whose function can impact hunger and feeding [67], and anxiety [68], as well as reduce the impacts of stress [69]. Dopa Decarboxylase (DDC), a gene which encodes the the enzyme that synthesizes norepinephrine (NE) from dopamine [70], is similarly more responsive in the female pituitary (FC_difference_= 4.76), as is Prolactin releasing hormone (PRLH) (ΔMedian= 1.082668). Actin gamma 2 (ACTG2) (ΔMedian= .75). The full table of differential expression results of genes more responsive to stress in the female pituitary as compared to the male pituitary is available at https://git.io/vDzQ1.

**Figure 6.**
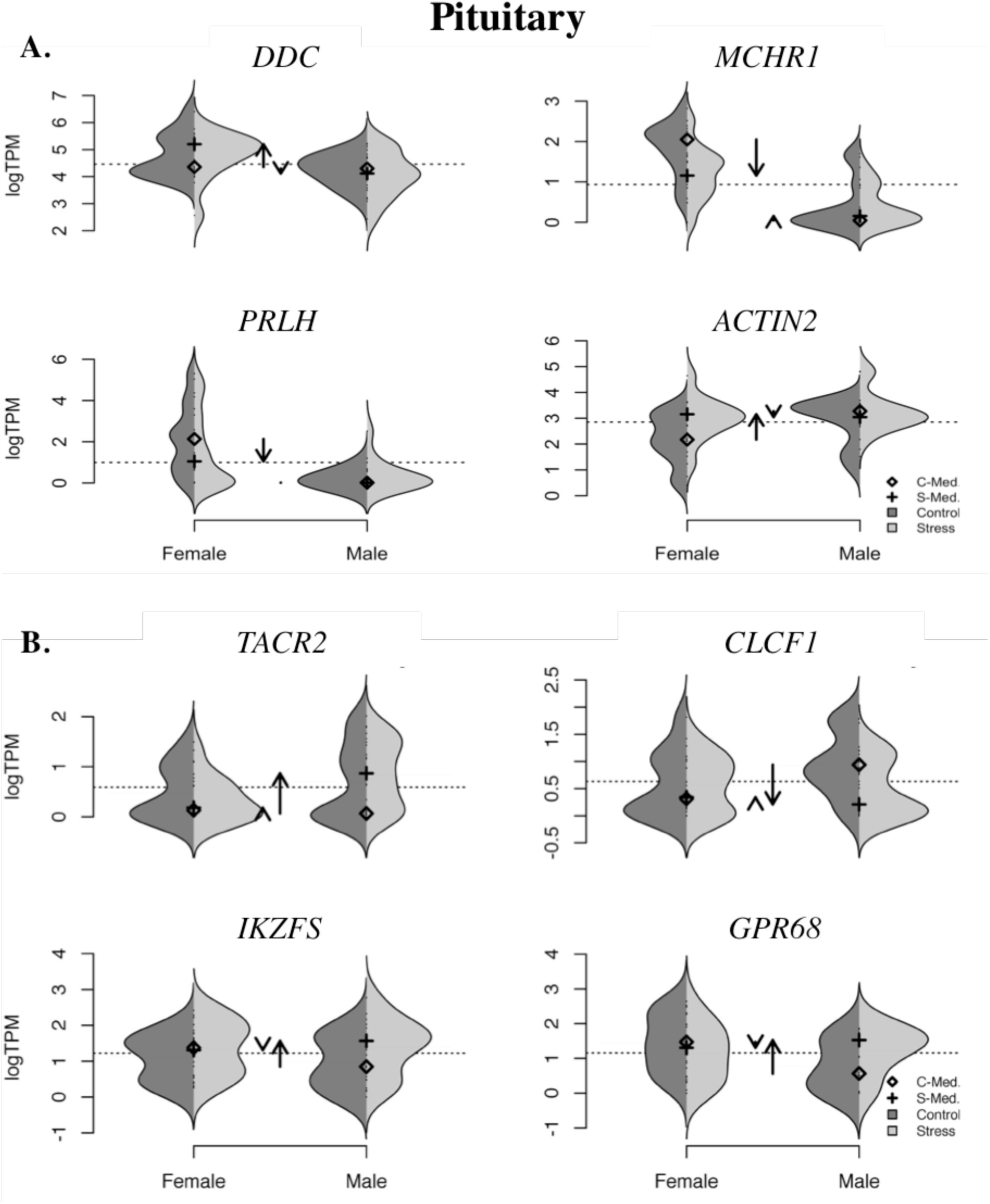
Violin plots depicting genes that were more responsive to stress in the a) female pituitary relative to the male pituitary, and b) male pituitary relative to the female pituitary. Plot titles are of the format Gene Name:Entrez ID:Tissue. The Y-axis indicates levels of gene expression as presented by taking the log of transcripts per million (logTPM). The shape of each half of the plot represents the kernel density estimation of the data, with darker grey indicative of data from control animals and lighter grey from stressed animals. The diamond symbol represents the median value of the control group while the plus sign represents the median value of the stressed group. Arrows represent the direction and amount of change in gene expression in the stressed group as compared to controls (up for upregulated, down for downregulated).

Genes more responsive to stress in the male pituitary relative to the female pituitary (examples in Fig. 6b) include tachykinin receptor 2 (TACR2, ΔMedian= -1.37), a gene known to have important reproductive correlates, specifically in loss of sexual cyclicity in females [71,72].[71,72] Genes that play key roles in the immune response, cardiotrophin-like cytokine factor 1 (CLCF1) [73,74]) and IKAROS family zinc finger 3 (IKZF3) [75,76] G protein coupled receptor (GPR68)(ΔMedian = 0.79). The full table of differential expression results of genes more responsive to stress in the male pituitary as compared to the female pituitary is available at https://git.io/vDzA8.

## DISCUSSION

By leveraging a highly-replicated and sex-balanced experimental approach to understand the transcriptomic effects of stress on the HPG axis, we provide an unparalleled glimpse into how males and females respond to stress. We exceeded replication standards for RNAseq experiments [77], while performing gene-level quantitation [78], which resulted in exceptionally robust estimates of gene expression across tissues and treatments. Our results provide evidence of sex-biased gene activity of a tissue system vital for vertebrate reproduction - the hypothalamus in the brain, the pituitary gland, and the gonads (testes and ovaries).

More genes in the female HPG axis were responsive to stress as compared to males. This differential expression of genes in response to stress was also mostly unique to each sex. Various factors could contribute to a sex-biased response to stress, though we were able to control for many of them with the rock dove model. For example, an uneven parental care strategy (one sex cares more than the other) and age have been known to influence the stress response [79–81]. However, both rock dove sexes offer significant offspring care, though they were not caring for offspring at the time of collection, and all birds were of similar age (2 years old). Males and females of this species are similar in other ways: they are physically monomorphic, socially and genetically monogamous [82], and we report that both sexes exhibit a similar increase in circulating glucocorticoids in response to restraint stress. This classic method of measuring the effects of stress on circulating glucocorticoid concentration would have suggested that rock dove males and females have a similar stress response. However, a deeper look at gene transcription in response to stress suggests otherwise. This begs the question as to whether a gene(s) is regulated in a sex-specific manner to produce a sex-specific physiological or behavioral result in the face of stress, such as female control over reproductive timing? Or, does expression resulting from stress lead to sexually monomorphic gene-mediated reproductive processes that converge to similar physiological and behavioral endpoints [83], such as the similar corticosterone response we observed? These data we provide can inform and promote multiple lines of further investigations to answer these questions, including hypothesis driven tests and manipulations of newly identified sex-biased and stress-responsive genes. Here, we propose three, non-mutually exclusive hypotheses to explain why more genes are responsive to stress in the female HPG axis as compared to their male counterparts: the Reproductive Cycle Hypothesis, the Reproductive Investment Hypothesis, and the Environmental Preparation Hypothesis.

Sexually mature females, unlike sexually mature males who maintain a relatively consistently functioning HPG axis, experience cycling of their reproductive hormones to facilitate ovulation and follicle growth; because of this, females may present a more complicated picture for understanding the effects of stress on the HPG axis, especially due to the potential for feedback mechanisms to change with the reproductive cycle [3,84]. The ovary was the site of the most differential expression in response to stress as compared to the hypothalamus, pituitary, and testes. Although we controlled for reproductive stage, selecting sexually mature birds that were not actively breeding, as well as specific ovary tissue type and amount sampled, we were unable to control for the specific stage of ovulation and follicle maturation at the point of sampling. Endocrine processes associated with these reproductive processes thus may influence how the HPG axis, particularly the ovary in this case, responds to environmental perturbations such as stress. This may be why we found a greater array of genes active in the female at both baseline sampling [24] and in response to stress. Alternatively, potential endocrine variation experienced by the female HPG axis might create “noise” and thus *decrease* our statistical ability to identify differentially expressed genes. Because we observed a significant increase, not decrease, in differentially expressed genes in the female HPG axis at baseline and in response to stress as compared to males, this is either not the case, or there is even more differential expression in female HPG tissue than we were able to statistically uncover. In either event, females are experiencing heightened HPG gene activity in response to stress, and this may be due to the physiological variation they experience over the course of their reproductive period.

Another potential hypothesis to explain why females are more responsive to stress than males at the level of their HPG transcriptome is an evolutionary one related to their reproductive investment. Though both males and females of this species are socially and genetically monogamous and offer biparental care [82], females generally invest more time and resources in making and maturing gametes and eggs [85,86]. Thus, because reproduction is arguably more energetically expensive for females during gamete and egg production, this may have supported the adaptation of a more stress-responsive reproductive axis to influence when and how females breed in response to the environment to optimize their lifetime reproductive success [87]. For example, prolactin is a hormone that is involved in a multitude of biological processes, but it has most notably been studied for its influential role in lactation and parental care in both birds and mammals. In this experiment, *Prolactin* in the hypothalamus and pituitary decreased in expression in response to stress in females but not males. Both male and female doves use prolactin to facilitate reproductive behaviors like nest building, lactation, and offspring care [88–90]. While males and females are responsive to stress at every level of their reproductive axis, a heightened responsiveness of reproductive substrates like prolactin by females could have evolved to support their need for increased sensitivity to the environment due to their higher level of reproductive investment. In this same vein, glucocorticoid receptor (GR), which binds the stress response hormone corticosterone, increased in expression in the female but not male hypothalamus in response to stress, suggesting a potential increase in sensitivity to the stress response by the female reproductive axis.

In another example, the use of RNAseq helped to uncover a less well-studied gene that could play a pivotal role in suppressing reproduction in the face of stress, *Calcitonin Gene-Related Peptide* (*CALCA*). *CALCA* was more responsive to stress in the female hypothalamus relative to the male hypothalamus. In another study, central administration of CALCA into the lateral cerebral ventricle of ovariectomized rats resulted in suppression of LH pulses, which was reversed by a CALCA receptor antagonist [61]. Stress-induced suppression of LH pulses was also blocked with a CALCA receptor antagonist [61]. These data suggest CALCA could be involved in stress-induced suppression of the reproductive axis. Here, we report that expression of the gene for the CALCA peptide is responsive to stress, more so in females than in males. However, *CALCA* gene expression decreased in response to stress. A decrease in expression might suggest a decrease in peptide production, which seems counterintuitive to the reported actions of CALCA in rats. However, the species, its physiology (for example, our birds were not ovariectomized), the time course of sampling, and the potential for physiological feedback must all be considered to gain a better picture of *CALCA* regulation. For now, its responsiveness to stress in females coupled with reports of its actions [61] suggest it may play an important role in regulating the reproductive system, and this may be related to reproductive investment.

A third hypothesis to explain increased female HPG genomic responsivity to stress is that information about the external environment experienced by the female could shape embryo, egg, and chick development, potentially priming offspring in a such a way as to increase their fitness in that stressful environment. For example, global warming is a type of stress that poses a threat to the survival of many species. Zebra finch parents acoustically signal high ambient temperatures to their egg-bound embryos, adaptively altering their behavior, growth, reproductive success, and thermal preferences as adults [91]. Indeed, maternal exposure to prenatal or postnatal stress can alter the stress response and behavior in offspring [92–94]. Thus, herein lies the potential for the environment experienced by the female to influence the development of young pre-egg lay. Female rock doves experienced a significant change of expression in stress- and reproduction-related genes in response to stress (Table 1) as well as in those related to immune function, growth, and other processes (eg. a gene ontology term enriched for genes more highly expressed in the ovary in response to stress was related to the inflammatory response). Resulting alterations to physiology and behavior could lead to maternal effects of epigenetic and genomic activity, setting into motion biological events that could prepare offspring for the environment in which they will soon face.

In summary, we report sex-specific changes in gene expression of the rock dove HPG axis in response to stress, with females exhibiting increased genetic responsiveness at all levels of their axis as compared to males. This phenomenon could be explained by the variation females experience in their reproductive cycle as well as by their evolutionary history, including parental investment and the potential for maternal effects to increase the reproductive success of offspring. These hypotheses are not mutually exclusive and inspire future investigations of genome to phenome causal effects. Presently, the data we report create a vital genomic foundation on which sex-specific reproductive dysfunction and adaptation in the face of stress can be further studied.

## ACKNOWLEDGEMENTS

We thank the many undergraduate and graduate students of the Calisi Lab for their assistance on this project, including their involvement in avian husbandry, conducting the experiment, and tissue collection. We also thank the UC Davis Environmental Endocrinology Group, comprised of the R. Calisi, T. Hahn, M. Ramenofsky, K. Ryan, and J. Wingfield Lab groups, for their discussion of this project. This work was funded by NSF IOS 1455960 (to RMC and MM).

## MATERIALS AND METHODS

### Animal Collection Methods

Birds were housed at the University of California, Davis, in large aviaries (5’x4’x7’), with 8 sexually reproductive adult pairs per aviary. Food and water were provided *ad libitum*. To control for reproductive stage and potential circadian rhythm confounds, we sampled males and females that were paired but lacked eggs or chicks between 0900-1100 (PST) following animal care and handling protocols (UC Davis IACUC permit #18895). Birds in the baseline, or “unstressed,” group were sampled within 5min of entering their cage. Birds in the stress treatment group were restrained in cloth bags for 30min prior to sampling. To sample tissue, birds were first anesthetized using isoflurane until unresponsive (<2 min), at which point they were decapitated. Trunk blood was collected to assay for plasma corticosterone concentrations. Brains, pituitaries, and gonads were then immediately extracted and placed on dry ice, then transferred to a -80 C freezer until further processing. Brains were sectioned coronally on a cryostat (Leica CM 1860) at 100*μ*M to best visualize and biopsy the hypothalamus. We used Karten and Hodos’ [95] stereotaxic atlas of the brain of the pigeon to locate the hypothalamus and collect it in its entirety. In brief, we, collected hypothalamic tissue beginning at the point of bifurcation of the tractus septomesencephalicus and ending afer the cerebellum was well apparent. Lateral septum tissue was included with the hypothalamus. We sequenced tissue from whole homogenized testes and ovaries, the latter comprised of tissue from the oviduct and ovarian follicles. Hypothalamic sections, pituitaries, and gonads were preserved in RNALater and shipped from the UC Davis to the University of New Hampshire for further processing. This technique to harvest hypothalamic, pituitary, and gonadal tissue from this species has been previously validated by our research group [24].

### Hormone assay

Fresh blood was centrifuged at 4200 rpms at 4C for 10min. Plasma was removed and stored at - 80C. We assayed plasma for corticosterone using radioimmunoassay (RIA), informed by a serial dilution conducted prior to the assay. A dilution of 1:20 was used in a commercially available Corticosterone RIA kit (MP Biomedicals, Orangeburg, NY) to determine corticosterone levels (ng/ml). The assay was validated for cross-reactivity with *C. livia* corticosterone and the limit of detection was estimated at 0.0385 ng/ml.

### Illumina Library Preparation and Sequencing

Tissues frozen in RNALater were thawed on ice in an RNAse-free work environment. Total RNA was extracted using a standard Trizol extraction protocol (Thermo Fisher Scientific, Waltham, MA). The quality of the resultant extracted total RNA was characterized using the Tapestation 2200 Instrument (Agilent, Santa Clara, CA), after which Illumina sequence libraries were prepared using the TruSeq RNA Stranded LT Kit (Illumina). The Tapestation 2200 Instrument was, again, used to determine the quality and concentration of these libraries. Each library was diluted to 2nM with sterile ddH2O, and pooled in a multiplexed library sample. The multiplexed library sample was then sent to the New York Genome Center for 125 base pair paired-end sequencing on a HiSeq 2500 platform.

### Transcriptome assembly evaluation and improvement

The previously constructed Rock Dove transcriptome version 1.0.3 assembly [24] was evaluated to ensure that transcripts expressed uniquely in the stress condition were included. To accomplish this, reads from the pituitary, hypothalamus, and gonads from one male and one female stressed individual were assembled following the Oyster River Protocol [96]. Unique transcripts contained in this assembly relative to the previously described assembly were identified via a BLAST procedure. Novel transcripts, presumably expressed uniquely in the stress condition were added to the existing assembly, thereby creating the Rock Dove version 1.1.0 transcriptome. This new assembly was evaluated for genic content via comparison with the BUSCO version 2.0 Aves database [97].

### Mapping and Global Analysis of Differential Gene Expression

After quality and adapter trimming to a Phred score =2, reads were quasimapped to the Rock Dove version 1.1.0 transcriptome after an index was prepared using Salmon 0.7.2 [98]. Rock dove transcript IDs were mapped to genes from *Gallus gallus* genome version 5, using BLAST [99]. All data were then imported into the R statistical package (version 3.3.0) [100] using tximport [101] for gene level evaluation of gene expression, which was calculated using edgeR (version 3.1.4) [102] following TMM normalization and correction for multiple hypothesis tests by setting the false discovery rate (FDR) to 1%. Gene ontology enrichment analysis was carried out using the Kolmogorov-Smirnov test [103] for significance in the R package, topGO [104]. A select set of genes found to be differentially expressed were plotted (*e.g.,* Figures 5 and 6) using the ‘beanplot’ package available at https://cran.r-project.org/package=beanplot.

### Candidate Gene Expression Evaluation

To evaluate a set of candidate genes, we selected *a priori* genes of interest based on their known involvement in stress and reproduction and associated behaviors (Table 1). Differences in gene expression were evaluated between these genes in the hypothalamic, pituitary, and gonadal tissues and between both sexes using a generalized linear model framework (expression ~ sex * tissue * treatment) with significance for all pairwise combinations of factors tested using the Bioconductor package lsmeans (https://cran.r-project.org/package=lsmeans) after correction using a dunnettx adjustment.

### Differential Response to Stress

In addition to understanding patterns of differential gene expression between stressed and control birds, genes were identified whose response to stress varied by sex. A distribution was generated corresponding to the absolute difference in median gene expression between treatment and sex. Genes located in the upper and lower 1% of this distribution were retained as significantly different in the response to stress.

